# Benzyl Cyanide Triggers A Regeneration Program in Arabidopsis

**DOI:** 10.64898/2026.07.20.738417

**Authors:** Julia Falińska, Barbara Wójcikowska, Marek Marzec

## Abstract

Phenylacetic acid (PAA) is a naturally occurring auxin whose biosynthesis and function during plant regeneration remain poorly understood. PAA may be synthesized from phenylalanine via the CYP79A2-dependent phenylacetaldoxime pathway, in which benzyl cyanide/phenylacetonitrile (BnCN/PAN) is a potential intermediate and substrate for NITRILASE (NIT) enzymes. Here, we investigated whether BnCN/PAN promotes somatic embryogenesis (SE) through NIT-dependent PAA biosynthesis. Low concentrations of BnCN/PAN stimulated somatic embryo formation in Arabidopsis thaliana, whereas exogenous PAA also promoted embryogenic induction. Transcriptome profiling revealed that BnCN/PAN upregulated genes associated with SE, including key embryogenic regulators and EMBRYO DEFECTIVE genes. RNA-seq data further indicated enhanced auxin signalling, which was independently confirmed using the pDR5::GUS reporter line. Inhibition of NIT activity by heatin reduced the embryogenic competence of BnCN/PAN-treated explants, supporting the involvement of NIT enzymes in this response. Collectively, our findings provide the first evidence that BnCN/PAN promotes embryogenic transition and suggest that NIT functions in SE extend beyond their proposed role in indole-3-acetic acid biosynthesis.

**Summary statement:** This study uncovers a previously unrecognized pathway regulating plant regeneration, providing new insights into how naturally occurring metabolites influence embryo formation.

## Introduction

Benzyl cyanide (BnCN), also known as phenylacetonitrile (PAN), is an aromatic nitrile that is essential for biochemistry (Miki and Asano, 2014), agriculture (Noge and Tamogami, 2013, Lee et al., 2025), and biotechnology (de Oliveira et al., 2013). BnCN/PAN chemical synthesis typically relies on Kolbe reaction between benzyl chloride and sodium cyanide or toluenes and cyanogen chloride (Chidambaram et al., 2007, Grimm and Menting, 1975). BnCN/PAN has been applied extensively in synthetic organic chemistry, where it serves as a versatile substrate or intermediate for a valuable chemicals synthesis used in manufacture of fragrances (ester compounds), textile industry (optical bleaches for fibers), pharmaceuticals (synthetic penicillins, barbiturates), and agrochemicals (insecticides) due to its reactivity and solubility characteristics (Miki and Asano, 2014, Colombo et al., 1984, Zhang et al., 2013, Pollak et al., 2000, Hou et al., 2015).

In living organisms, endogenously synthesized BnCN/PAN, has emerged as an important metabolite with diverse biological properties and physiological roles in plants (Yamaguchi et al., 2023, Wheeler, 1977) and animals (Wei et al., 2019). In the plant sciences, nitriles such as BnCN/PAN have gained attention for their biological activities and roles as signaling molecules, metabolic intermediates, and defense compounds. When BnCN/PAN applied exogenously to plants induces dose-dependent morphological and developmental changes that resemble classic auxin overproduction phenotypes (Wheeler, 1977, Wheeler et al., 1980). These include significant reductions in primary root length and total biomass, elongation of hypocotyls, epinastic cotyledon formation, and increased adventitious root emergence - traits that are consistent with elevated auxin signaling and homeostasis perturbation (Urbancsok et al., 2018). In plants, the physiological and developmental impacts of endogenously synthesized and exogenously applied BnCN/PAN are mediated through enzymatic conversion to active metabolite i.e. phenylacetic acid (PAA), a naturally occurring auxin. PAA is widely present in plant species, however its synthesis, conjugation and action are still not fully understood, and during the course of investigation (Perez et al., 2023b, Hladík et al., 2025, Prinsen, 2025). It is known that PAA is not polarly transported in plants, showed higher accumulation than IAA, and PAA level depends on the plant species or the studied organ (Perez et al., 2023b, Sugawara et al., 2015, LudwigCMüller and Cohen, 2002, Prochazka and Borkovec, 1984, Morris and Johnson, 1987, Cook, 2019). PAA has less auxin activity than IAA but regulates the expression of auxin-responsive genes with the contribution of auxin receptors (Shimizu-Mitao and Kakimoto, 2014, Sugawara et al., 2015, Cook et al., 2021, Smit and Went, 1935, Lee et al., 2014, Napier and Venis, 1990). PAA promotes root growth, lateral root formation, elongation of coleoptiles and internodes in various species (Cook, 2019, Perez et al., 2023b), as well as poses antimicrobial properties (Pan et al., 2022, Kim et al., 2004, Mao et al., 2006, Zhang et al., 2022, Small and Morris, 1990). Parallel to IAA synthesis pathways, also appear to be the case for the biosynthesis of PAA, but the enzymes involved in the respective pathways might differ (Cook et al., 2016). PAA is synthesized from the phenylalanine (Phe) precursor through TAA-dependent phenyl-pyruvate (PPyA), CYP-dependent phenylacetaldoxime (PAOx) pathways, but also might be potentially synthesized from benzyl glucosinolates (BGs) (LudwigCMüller and Cohen, 2002, Prinsen, 2025, Cook et al., 2016). PPyA pathway involves YUCCAs, a flavin monooxygenase having a key role in auxin biosynthesis (Dai et al., 2013, Cook et al., 2016, Sugawara et al., 2015). Different CYP enzymes were identified to be involved in PAOx synthesis i.e.: CYP79A2, CYP79A61, CYP79B2, CYP79B3 in *Zea mays, Sorghum bicolor* and *Arabidopsis thaliana* (Aoi et al., 2020, Perez et al., 2021, Perez et al., 2023a, Irmisch et al., 2015). PAOx could be converted to BnCN/PAN or phenylacetamide (PAM) and then to PAA. Crucially, genetic and biochemical evidence underscores the role of plant NITRILASES in mediating BnCN/PAN’s effects through enzymatic conversion to PAA. Nitrilases are a class of hydrolytic enzymes that process organic nitriles into their corresponding carboxylic acids and ammonia. Studies in the *Arabidopsis thaliana* have provided pivotal insights into the auxin-like activities of BnCN/PAN and the enzymatic processes that mediate its potential conversion *in planta*. In *Arabidopsis thaliana* the NIT1-NIT3 subgroup exhibits catalytic activity toward BnCN/PAN, and loss-of-function mutants in these genes demonstrate altered sensitivity to exogenous BnCN/PAN, exhibiting mitigated growth responses compared to wild-type controls. In details, the single mutants *Atnit1* and *Atnit2* were less sensitive to BnCN/PAN than the wild-type, with *Atnit2* being less resistant than *Atnit1*, probably because *AtNIT2* exhibits tissue-specific expression and is not active in seedlings. Moreover, *Arabidopsis thaliana* transgenic line *nit2*-RNAi line with the silencing of all *NITs* show resistance to a broad range of BnCN/PAN concentrations, reinforcing the idea that NIT1-NIT3-mediated conversion is necessary for BnCN/PAN’s bioactivity. In contrast, direct application of PAA to wild-type and *Atnit1*, *Atnit2*, *nit2*-RNAi mutants of *Arabidopsis thaliana* plants produces robust auxin phenotypes, supporting the model that the biological effects of BnCN/PAN derive principally from its *in planta* conversion to PAA. So, the auxin-like effects triggered by BnCN/PAN on *Arabidopsis thaliana* are possibly connected with its NIT1-NIT3-mediated conversion to the natural PAA (Urbancsok et al., 2018). Among plant secondary metabolites, nitriles are most commonly linked to the glucosinolate-myrosinase system found in *Brassicales* species. In this system, the enzymatic breakdown of glucosinolates, such as benzyl glucosinolate may lead to the formation of various hydrolysis products, including BnCN/PAN compounds.

BnCN/PAN compounds influence plant-herbivore interactions also and can act as chemical defenses. Interestingly, the *PtNIT1*, *PtNIT2, PtNIT3* genes are upregulated in *Populus trichocarpa* following herbivory events (Günther et al., 2018). It was shown that PAN/BnCN accumulates in herbivore-damaged tissue in *Populus trichocarpa* and *Fallopia sachalinensis* (Günther et al., 2018, Noge and Tamogami, 2013, Yamaguchi et al., 2016, Noge et al., 2011). Additionally, *Camellia sinensis* is also known to emit BnCN/PAN in response to postharvest stress (Liao et al., 2020). PAN/BnCN might also be catabolized to PAA as treatment of *Populus trichocarpa* leaves with (α-^13^C)PAN/BnCN resulted in the formation of (α-^13^C)PAA. Additionally, it has been shown that BnCN/PAN is converted to PAA *in vitro* and is one of the substrates for the recombinant NIT1 enzyme in *Populus trichocarpa*, phylogenetically distinct from the *Arabidopsis thaliana* NIT1, NIT2, and NIT3, and is related to the NIT4-type NITs (Günther et al., 2018, Aoi et al., 2020). Summarizing, BnCN/PAN, itself functions as a plant defense compound but becomes phytotoxic at high concentrations, and to avoid toxic PAN effects, it may be metabolized in plant cells to PAA (Günther et al., 2018).

Our research showed that in *Arabidopsis thaliana in vitro* culture, BnCN/PAN activates the embryogenic program, highlighting its potential role as a regulatory metabolite rather than a passive intermediate. We propose that the conversion of BnCN/PAN to PAA is potentially mediated by NITRILASES, as pharmacological inhibition of NIT activity in the presence of BnCN/PAN markedly reduces somatic embryogenesis (SE) efficiency. These results provide functional evidence for a link between nitrilase-dependent metabolism and embryogenic competence. Notably, we report here for the first time transcriptomic reprogramming in plant cells exposed to BnCN/PAN, revealing extensive changes associated with activation of the embryogenic pathway.

A comprehensive dissection of BnCN/PAN action will not only deepen our understanding of auxin biology and the integration of secondary metabolism with developmental plasticity, but may also open new avenues for metabolic engineering and the rational optimization of plant tissue culture systems.

## Results

### BnCN/PAN turns on the embryogenic program

To assess the ability of BnCN/PAN, a potential precursor utilized by NITRILASES (NITs) for phenylacetic acid (PAA) biosynthesis, to induce SE process, the immature zygotic embryos of *Arabidopsis thaliana* were treated with a range of BnCN/PAN concentrations. BnCN/PAN efficiently activated the embryogenic program at concentrations of 25-500 µM (Fig. 1). As a control, embryos were cultured on E0 medium lacking BnCN/PAN, where they predominantly converted into seedlings (Fig. 1C, C1). The response of the explants varied in a concentration-dependent manner following BnCN treatment. Explants responded to BnCN/PAN treatment with an SE efficiency of approximately 30% (Fig. 1A), producing on average 1.3 somatic embryos per explant (Fig. 1B). However, increasing concentrations of BnCN/PAN resulted in a progressive decline in both the efficiency and productivity of SE process. Upon BnCN/PAN treatment, somatic embryos regenerated from cells on the adaxial side of the cotyledons, and SE induction was accompanied by the formation of callus tissue and roots (Fig. 1D, D1-K, K1). For subsequent experiments, a concentration of 50 µM BnCN/PAN was selected, as it elicited the most stable and uniform response among explants, with 40% efficiency and 1.5 productivity of SE. A concentration of 600 µM BnCN completely inhibited SE process, explants failed to form even callus tissue (Fig. 1L, L1), indicating that this concentration is toxic to explant cells. Thus, BnCN/PAN can be considered an inducer of SE in *Arabidopsis thaliana*, although it is markedly less potent than 2,4-dichlorophenoxyacetic acid (2,4-D), which induces SE with an efficiency approaching 90% and a productivity of approximately four embryos per explant (Gliwicka et al., 2013).

**Figure 1.**
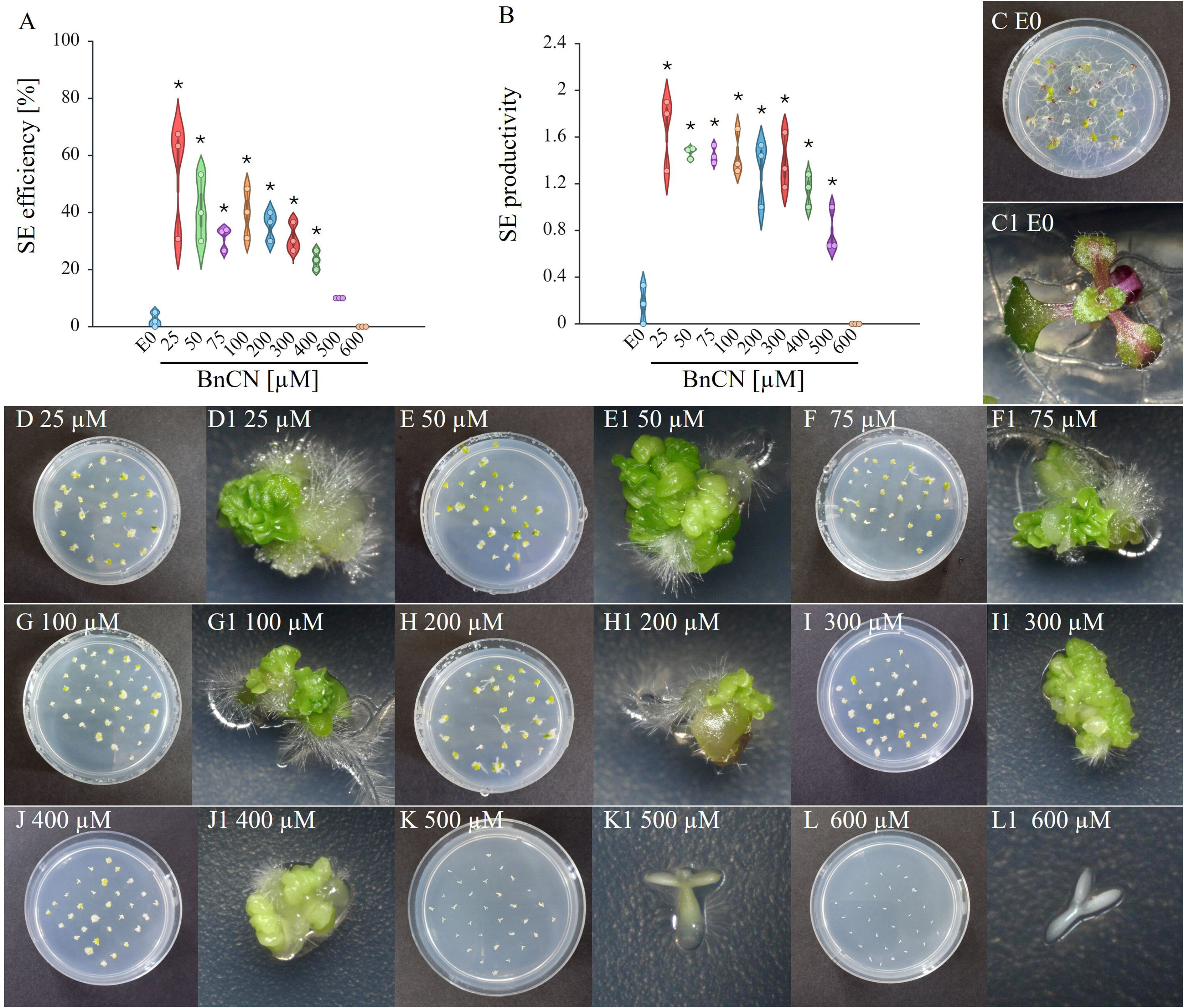
BnCN/PAN treatment induces embryogenic transition in immature zygotic embryo *in vitro* culture. The embryogenic capacity: an efficiency (A), and a productivity (B) of the Col-0 on the control E0 medium (C,C1) and EB medium supplemented with increasing BnCN/PAN concentration: 25 (D,D1), 50 (E,E1), 75 (F,F1), 100 (G,G1), 200 (H,H1), 300 (I,I1), 400 (J,J1), 500 (K,K1), 600 (L,L1) µM. SE capacity was evaluated in 21-day-old cultures. Values significantly different from the control culture (E0) are marked with asterisks (*). A two-way ANOVA analysis (*p* < 0.05) followed by Tukey’s HSD was used to determine any values that were significantly different (\**p* < 0.05) (n ≥ 3; means ± SD are given). The 6 cm plate (C-L), and explant images (C1-L1).

### PAA, product of NITs activity, can induce somatic embryogenesis process

We hypothesized that the role of NITs during the induction of SE may be linked to the biosynthesis of PAA from its precursor BnCN/PAN. To test this hypothesis, it was essential to determine whether PAA itself, as the direct product of potential NITs enzymatic activity, possesses intrinsic SE-inducing capacity. To this end, explants were treated with a range of PAA concentrations (5-150 µM; Fig. 2). PAA concentrations up to 75 µM supported the regeneration of somatic embryos (Fig. 2D, D1-I, I1), whereas higher concentrations resulted in a complete inhibition of SE (Fig. 2J, J1-M, M1). The optimal concentration, at which explants regenerated somatic embryos with the highest efficiency (nearly 100%) and productivity (approximately two embryos per explant), was 40 µM PAA (Fig. 2H, H1). These findings demonstrate that both the PAA precursor BnCN/PAN and PAA itself are capable of activating the embryogenic program. Although PAA induces SE with an efficiency comparable to that of 2,4-D (Gliwicka et al., 2013), its embryogenic productivity is approximately twofold lower.

**Figure 2.**
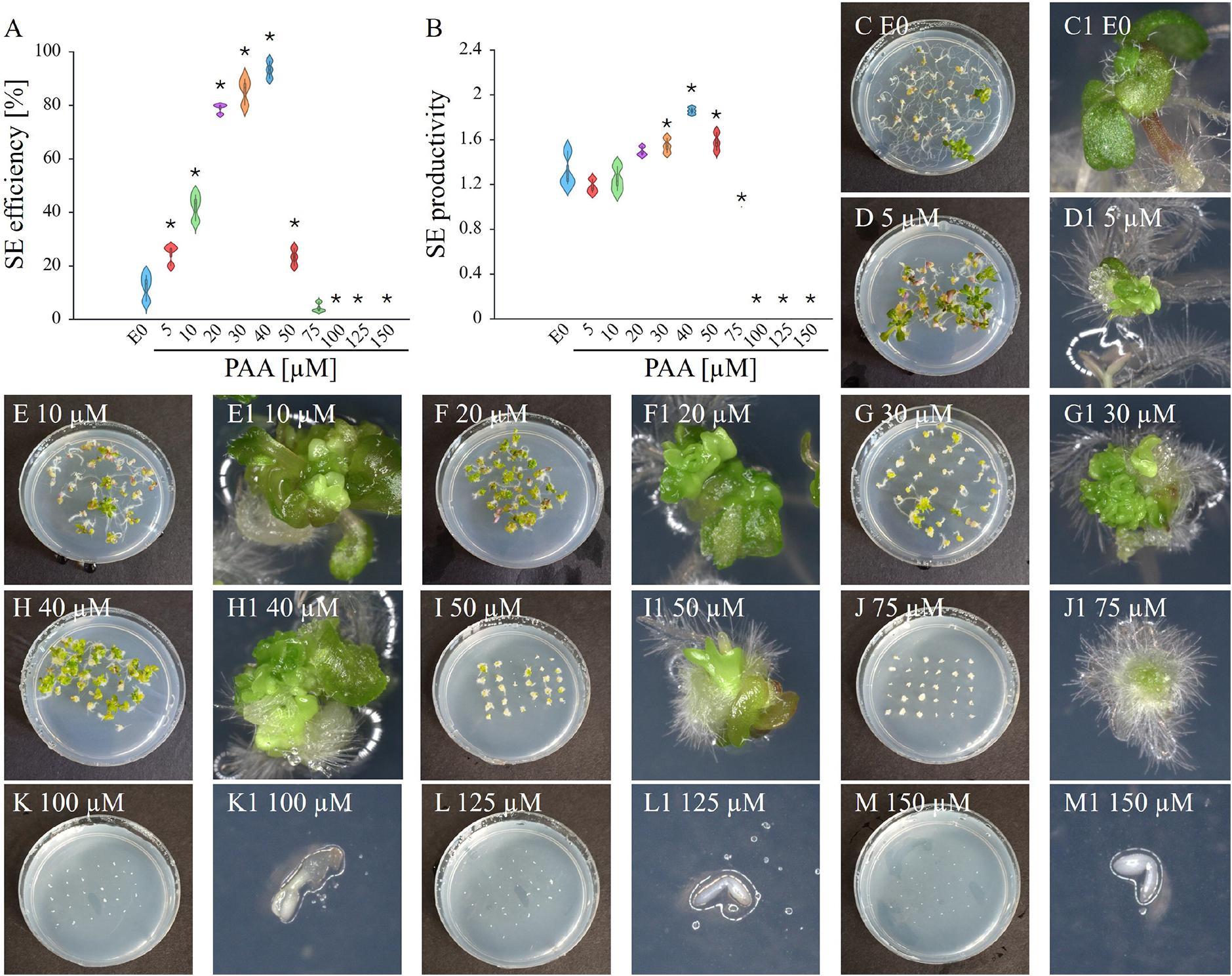
PAA treatment induces embryogenic transition in immature zygotic embryo *in vitro* culture. The embryogenic capacity: an efficiency (A), and a productivity (B) of the Col-0 on the control E0 medium (C,C1) and medium supplemented with increasing PAA concentration: 5 (D,D1), 10 (E,E1), 20 (F,F1), 30 (G,G1), 40 (H,H1), 50 (I,I1), 75 (J,J1), 100 (K,K1), 125 (L,L1), 150 µM (L,L1). SE capacity was evaluated in 21-day-old cultures. Values significantly different from the control culture (E0) are marked with asterisks (*). A two-way ANOVA analysis (*p* < 0.05) followed by Tukey’s HSD was used to determine any values that were significantly different (\**p* < 0.05) (n ≥ 3; means ± SD are given). The 6 cm plate images (C-M) and explant images (C1-M1).

### Evaluation of somatic embryogenesis competence in *nit* mutant lines

The impact of mutations in *NIT* genes on BnCN/PAN-dependent SE was assessed using the *nit1-3*, *nit2*, *nit3*, *nit4* mutants. The results indicate that only the *nit1*-*3* mutant exhibited a 35.7% reduction in SE efficiency compared to the wild-type (Fig. 3A, C-D). All other analyzed mutants regenerated somatic embryos at a frequency comparable to wild-type Col-0 (Fig. 3A, C, E-G). Similarly, all analyzed mutants displayed productivity (approximately 1.8 somatic embryos per explant) levels equivalent to the wild-type Col-0 (Fig. 3B, C1-G1). These findings suggest that the investigated *NIT* genes may act redundantly, which could explain the absence of observable effects in *nit2*, *nit3*, *nit4* single-gene mutants.

**Figure 3.**
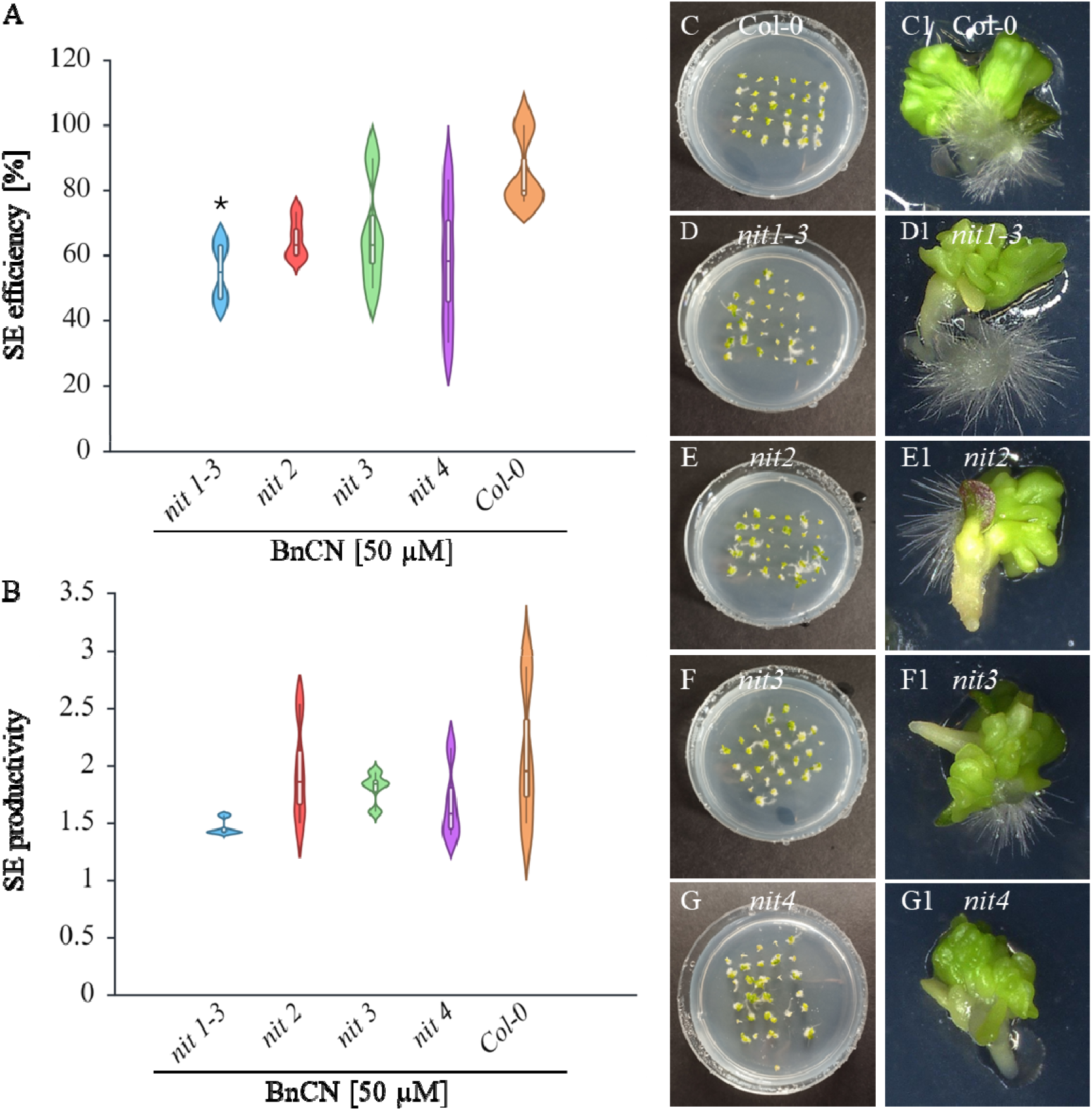
The embryogenic response of *nit* (*nit1-4*) mutants. Analysis of the effectivity (A) and productivity (B) of SE in the culture of wild-type (Col-0) (C,C1), *nit1-3* (D,D1), *nit2* (E,E1), *nit3* (F,F1), *nit4* (G,G1) mutants explants cultured on EB medium supplemented with 50 µM BnCN/PAN. SE capacity was evaluated in 21-day-old cultures. (*) significant differences between *nit* mutants and Col-0 (*p*C<C0.05, Student’s t test). The 6 cm plate images (C-G) and explant images (C1-G1).

### The inhibition of NITs by heatin negatively affects the ability of explants to undergo embryogenic transition under BnCN/PAN and 2,4-D treatment

Given the potential functional redundancy among NITs, we employed the established NIT inhibitor heatin (van der Woude et al., 2021) to pharmacologically suppress NIT activity. Simultaneous treatment of explants with the NIT inhibitor heatin (15 µM) and SE inducers-BnCN/PAN (50 µM) or 2,4-D (5 µM), led to a pronounced reduction in both the efficiency and productivity of SE (Fig. 4). Treatment of explants with heatin under BnCN/PAN-dependent SE induction resulted in an approximately 2.5-fold reduction in the proportion of explants forming somatic embryos. The obtained results may indicate that BnCN/PAN is a substrate for NITs, which convert it into the naturally occurring auxin PAA, resulting in the activation of the regeneration program.

**Figure 4.**
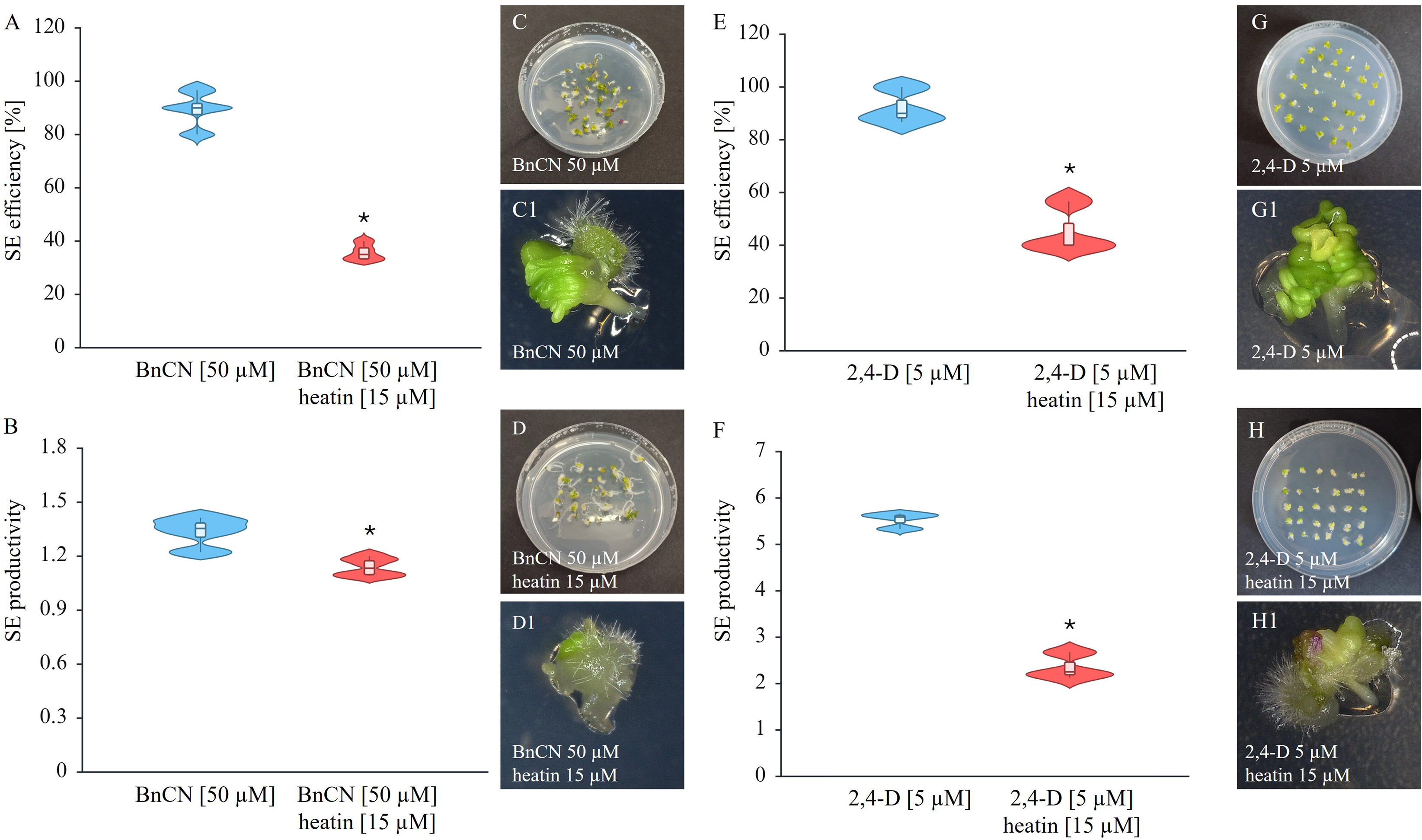
Inhibition of NITs by heatin reduces embryogenic response in *in vitro* culture of Col-0 explants. Analysis of the effectivity (A, E) and productivity (B, F) of SE in the culture of wild-type (Col-0) (C,C1), explants cultured on EB (A-D) or EA (E-H) media supplemented with 50 µM BnCN/PAN or 5 µM 2,4-D, respectively and additionally treated with 15 µM heatin (NIT-inhibitor). SE capacity was evaluated in 21-day-old cultures. (*) significant differences between control (EB, EA) media and EB, EA media supplemented with heatin (*p*C<C0.05, Student’s t test). The 6 cm plate images (C-G) and explant images (C1-H1).

### BnCN/PAN treatment caused robust changes in transcriptome

Next, the RNA-seq-generated transcriptomes of the BnCN/PAN-induced cultures were analyzed to get insights into the plausible BnCN/PAN-NITs-PAA mechanism of SE induction. The IZE explants at the cotyledonary stage of development were cultured on two media: EB supplemented with BnCN/PAN (50 µM), and E0 control, SE-inducer free medium. In EB medium, the first somatic embryos became visible on the adaxial side of the IZE cotyledons and shoot apical meristem on the 8-10^th^ day. In contrast, the explants that were cultured on a non-embryogenic medium (E0) developed into seedlings. The EB-and E0-cultured explant tissues were sampled for RNA-seq analysis at 10^th^ day of SE induction. Significantly regulated genes were selected under a threshold of FDR ≤ 0.05 and |log2FC| ≥ 1. Using these criteria list of differentially expressed genes (DEGs) in EB culture was generated in comparison to E0 culture (Additional file 1). Volcano plots were used to illustrate the distribution of the log2FC and FDR within the analyzed gene sets (Fig. 5A). Among all 26 435 analyzed genes 25% (6 853) were classified as differentially expressed (Additional file 2). Moreover, downregulated transcripts were found more frequently than upregulated under SE BnCN/PAN induction treatment. Accordingly, 75.8% (5 195) and 24.2% (1 658) of DEGs showed a significantly decreased (Additional file 3) or increased (Additional file 4) expression in response to BnCN/PAN (DEGs-EB/E0) treatment, respectively (Fig. 5A). Noteworthy, at least one-fifth of the downregulated DEGs (1 009) in embryogenic cultures showed a high decrease in transcript abundance by at least fivefold in EB treatment. Substantially less frequent were genes (85) of highly increased expression (FC≥5) in EB culture. Cluster analysis indicated that most DEGs were expressed during SE induction at lower levels than during seedling development (Fig. 5B). As displayed in Figure 5C, subclusters 1, 2, and 4 contained 109, 5 080, and 6 genes, respectively (Additional files 5-7), all of which were downregulated during BnCN/PAN dependent SE induction relative to control condition (E0). Conversely, subcluster 3 (1 658) (Additional file 8) exhibited upregulated expression in during embryogenic transition relative to seedling development and among them positive regulators of SE can we identified. Gene Ontology (GO) enrichment analysis of DEGs revealed that BnCN/PAN treatment triggers a pronounced reprogramming of cellular processes. Upregulated genes in BnCN/PAN-treated explants were significantly enriched in pathways associated with cell cycle progression, cell division, and cytoskeleton organization, together with processes related to DNA alkylation and methylation, including RNA-directed DNA methylation (RdDM), a key mechanism of transcriptional gene silencing (Fig. 5D). Consistently, KEGG analysis indicated activation of plant hormone signal transduction, RNA polymerase activity, motor protein function, DNA repair pathways, and central carbon and nitrogen metabolism, including lipid, amino acid, sugar, and starch metabolic processes (Fig. 5E). In contrast, downregulated genes were significantly enriched in functional categories associated with secondary metabolism, stress responses, and photosynthesis (Fig. 5F). These included responses to wounding, hypoxia, and toxic compounds, reflecting a general attenuation of stress-responsive programs. KEGG pathway analysis further revealed suppression of phenylpropanoid biosynthesis, along with plant–pathogen interaction, MAPK signaling, tryptophan metabolism, glutathione metabolism, and pathways involved in nitrogen, starch and sucrose metabolism, as well as cutin, suberin, and wax biosynthesis (Fig. 5G).

**Figure 5.**
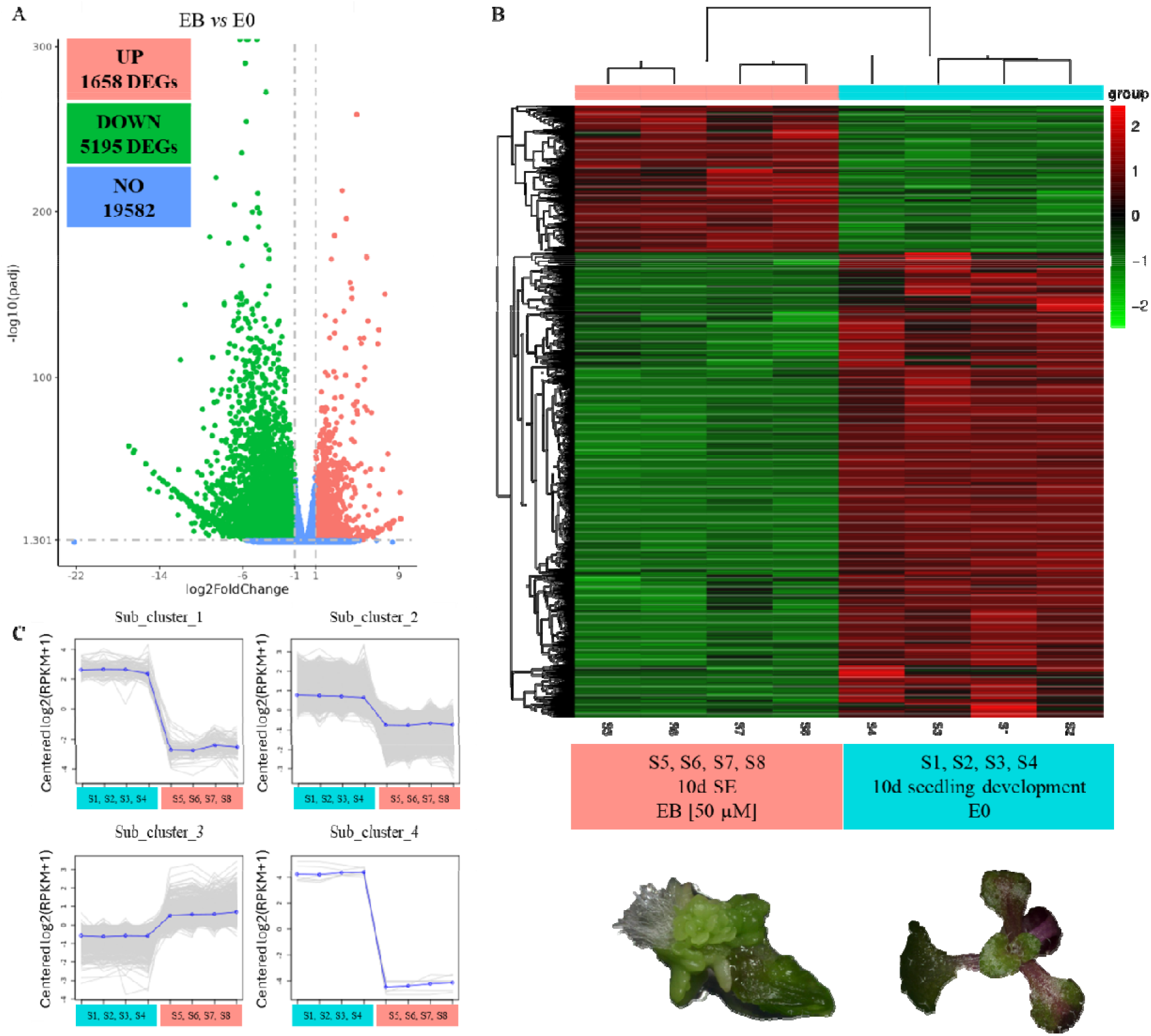
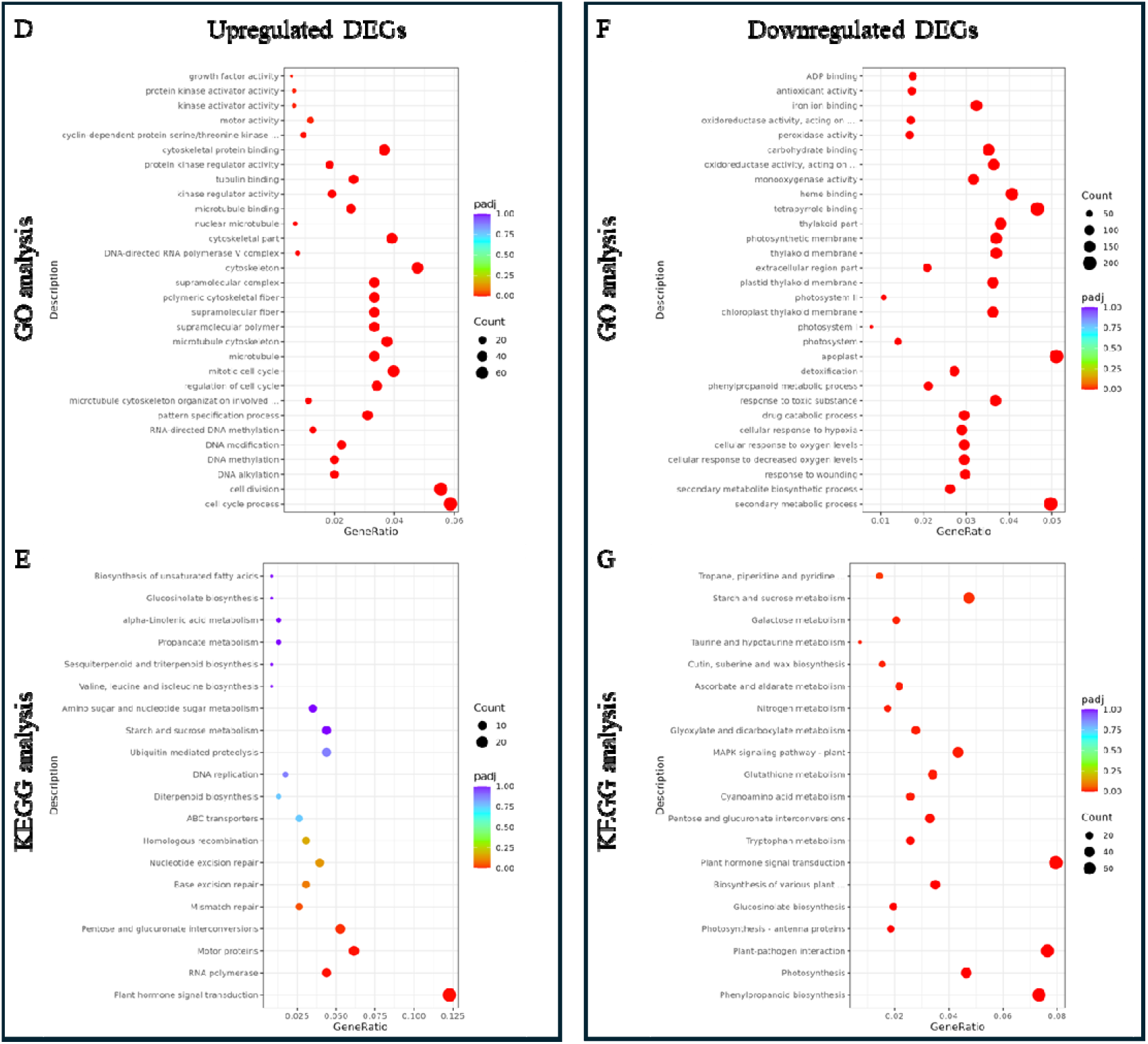
Gene expression profile and clustering analysis based on RNA sequencing analysis. Volcano plot of differentially expressed genes (DEGs) during the embryogenic transition induced by BnCN treatment on the 10^th^ day of *in vitro* culture. (A). Cluster analysis of DEGs in heat map (B). The color represents the transcriptional abundance of the DEGs. Red and green represent high and low expression levels, respectively. (C) K-means clustering of gene expression trends. The gray line represents the expression pattern of genes in each cluster, and the blue line represents the average expression of all genes in the cluster. Plant Ontology enrichment analysis of up-(D,E) and downregulated (F,G) DEGs during BnCN/PAN-dependent SE induction. S1-S4; S5-S8 - biological replicate.

### Convergent transcriptomic responses during somatic embryogenesis induction by different chemicals

To identify transcriptional programs associated with SE process irrespective of the inducing stimulus, we compared DEGs detected in explants treated with three mechanistically distinct SE inducers: the synthetic auxin 2,4-D, the histone deacetylase inhibitor trichostatin A (TSA), and BnCN/PAN. We compared publicly available datasets of DEGs at 10^th^ day of culture during SE process induced by 2,4-D and trichostatin A (TSA) (Wójcikowska et al., 2024), alongside DEGs identified here under BnCN/PAN-induced SE. Despite their different modes of action, all treatments triggered extensive and overlapping transcriptional responses. This analysis demonstrated that approximately 80.4% of DEGs (5 511 DEGs) following BnCN/PAN treatment are likewise deregulated during SE induced by other chemical compounds, including 2,4-D and TSA (Fig. 6A). Among DEGs, 74.7% were upregulated (1 239 DEGs) (Fig. 6B), whereas 78.2% (4 063 DEGs) were downregulated, and shared across SE-inducing treatments (Fig. 6C). The most prominent feature of the analysis was the identification of 3 659 DEGs shared among all three induction systems. Collectively, these results define a core set of 3 659 genes consistently deregulated during SE induction irrespective of the inductive stimulus and identify them as a strong candidate repository of regulators governing the transition from somatic to embryogenic cell fate. We hypothesize that genes critically involved in SE process are represented within a set of 3 659 genes differentially regulated independently of the SE-inducing treatment. Within this group, 756 genes exhibited increased expression and may represent putative positive regulators of SE, whereas 2 766 genes showed decreased expression and may function as potential negative regulators of the embryogenic transition (Additional file 9). This common gene set represented the largest overlap detected and indicates the existence of a conserved transcriptional framework underlying the acquisition of embryogenic competence. The substantial convergence of transcriptional responses suggests that distinct inductive signals ultimately channel cellular reprogramming through a common regulatory network. Thus, while auxin signaling, chromatin remodeling and BnCN/PAN-mediated responses may initiate SE through different upstream mechanisms, they appear to converge on a shared molecular program associated with embryogenic transition. Notably, the number of genes common to all treatments exceeded the number of treatment-specific DEGs identified for BnCN/PAN (1 342 genes), TSA (2 290 genes) and approached the size of the 2,4-D-specific fraction (3 287 genes). This observation highlights the predominance of a conserved transcriptional response over inducer-specific effects and suggests that the shared gene set is enriched in core regulators of embryogenic reprogramming. At the same time, each treatment elicited a substantial number of unique transcriptional changes, indicating that distinct molecular pathways contribute to the establishment of embryogenic competence. Among the analyzed treatments, 2,4-D induced the largest treatment-specific transcriptional response (3 287), consistent with the central role of auxin signaling in developmental reprogramming and the extensive remodeling of gene expression associated with auxin-dependent SE induction. TSA treatment also resulted in a large group of unique DEGs (2 290), supporting the notion that chromatin remodeling constitutes an important regulatory layer during SE induction. The transcriptional changes specific to TSA likely reflect the widespread derepression of genomic regions resulting from increased histone acetylation and altered chromatin accessibility. In contrast, BnCN/PAN generated the smallest set of treatment-specific genes (1 342) while exhibiting extensive overlap with the transcriptional responses induced by both TSA and 2,4-D. This pattern suggests that BnCN/PAN activates a more focused subset of pathways closely associated with embryogenic reprogramming rather than eliciting broad transcriptomic perturbations. Pairwise comparisons further revealed substantial similarities between auxin-and chromatin-mediated induction pathways. The overlap between 2,4-D and TSA comprised 2 774 genes, considerably exceeding the overlap observed between BnCN and 2,4-D (1 327 genes) or between BnCN/PAN and TSA (525 genes). These findings indicate that auxin signaling and chromatin remodeling engage highly interconnected regulatory networks during SE induction and support a model in which epigenetic reconfiguration represents an integral component of auxin-driven developmental reprogramming.

**Figure 6.**
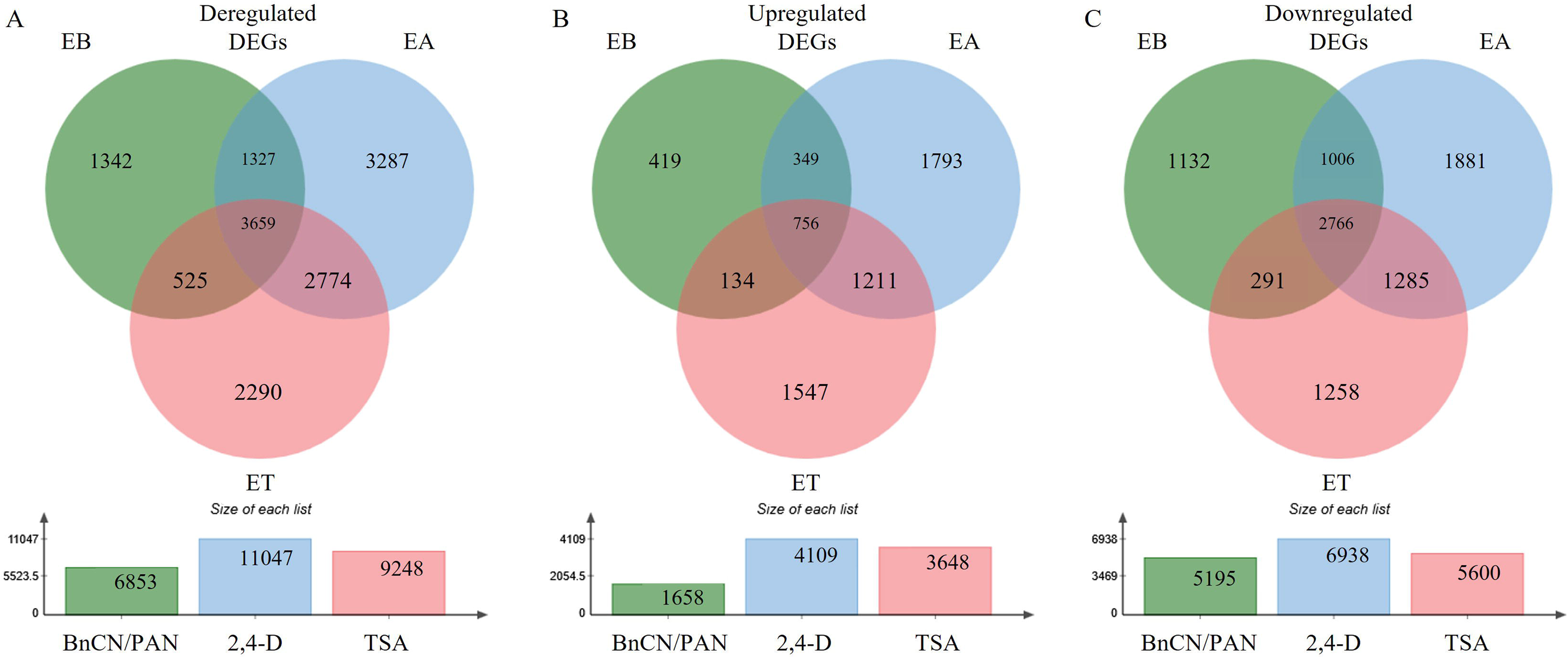
Transcriptomic response to BnCN/PAN treatment in Col-0 10^th^ day old *in vitro* embryogenic culture. The general number of DEGs (A), the number of upregulated (B), and downregulated (C) DEGs identified in embryogenic culture of Col-0 after BnCN/PAN (EB medium), 2,4-D (EA medium) and TSA (ET medium) treatment. Shared and treatment-specific DEGs in response to BnCN/2,4-D/TSA treatments were determined by comparing transcriptomic profiles with each other.

### Inducer-specific *NIT* genes expression during somatic embryogenesis

Analysis of *NIT* genes expression during SE revealed that BnCN/PAN treatment of explants led to increased expression of *NIT2* and *NIT4*, consistent with their upregulation during TSA-and 2,4-D-induced SE (Fig. 7A) (Wójcikowska et al., 2024). In contrast, *NIT3* displayed reduced expression, in agreement with patterns observed under both TSA and 2,4-D treatments. Notably, BnCN/PAN did not induce *NIT1* expression, whereas its regulation during SE was dependent on inducer, being upregulated under TSA-dependent SE induction and downregulated during 2,4-D-mediated SE induction (Fig. 7A). BnCN/PAN-mediated SE induction elicits *NIT* genes expression patterns largely consistent with those observed under TSA and 2,4-D treatments (Wójcikowska et al., 2024), indicating a shared regulatory framework, while the distinct, inducer-dependent regulation of *NIT1* highlights specific differences between these pathways.

**Figure 7.**
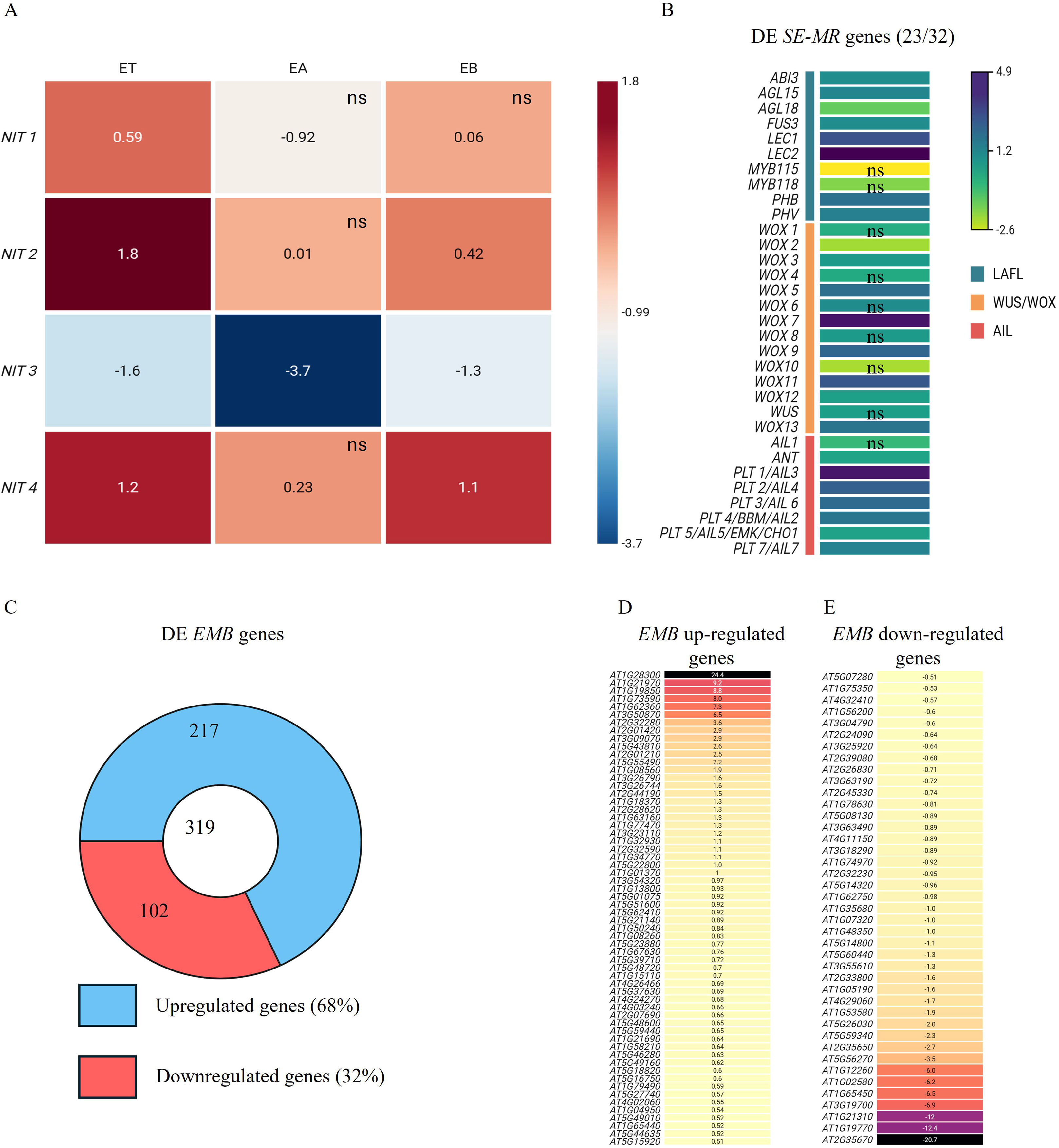
Transcriptional dynamics of genes encoding NITRILASEs (NITs), somatic embryogenesis master regulators (SE-MRs) and EMBRYO-DEFECTIVE (EMB) proteins during SE induction. Differential expression of *NIT1*–*NIT4* in explants cultured for 10 days on embryogenic media (ET, EA and EB). Values indicate relative transcript abundance (log2FC) in ET, EA and EB compared with E0 (A). Heatmap showing the expression profiles of transcription factors acting as key regulators of somatic embryogenesis (B). Number of differentially expressed *EMB* genes (C), including genes exhibiting increased (D) or decreased (E) transcript abundance. (ns) no significant differences in gene expression between E0 and ET, EA or EB explants at the corresponding developmental stage.

### BnCN/PAN activates a number of genes with confirmed and documented role in the embryogenic processes, including SE master regulators and *EMB* genes

We found most of the central SE-regulators, including *LEAFY COTYLEDON 1, 2* (*LEC1, 2*)*, FUSCA3* (*FUS3*)*, ABSCISIC ACID INSENSITIVE* 3 (*ABI3*)*, PHABULOSA* (*PHB*)*, PHAVOLUTA* (*PHV*)*, AGAMOUS-LIKE 15* (*AGL15*), and most of the *WUSCHEL-RELATED HOMEOBOX* (*WOX2*, 3, *5*, *7*, *9*, *11, 13*) and *PLETHORA* (*PLT*) gene family members (*AINTEGUMENTA*-*ANT*, *PLT1-PLT5*, *PLT7*) upregulated in response to BnCN/PAN treatment (Fig. 7B; Additional file 10). A particularly high increase of BnCN/PAN-dependent expression (FC from 20 to 24) showed *LEC2, WOX7,* and *PLT1/AIL3.* Among 510 identified *EMBRYO-DEFECTIVE* (*EMB*) genes (Meinke, 2020), 64% (319) were differentially expressed after BnCN/PAN treatment, and among them 68% (217) and 32% (102) were up-or down-regulated, respectively (Fig. 7C-E; Additional file 10).

### Increased auxin response accompanies the BnCN/PAN treatment

RNA-seq analysis revealed a profound and highly coordinated reprogramming of the auxin signaling network mediated by *AUXIN RESPONSE FACTORs* (*ARFs*) and *AUXIN/INDOLE-3-ACETIC ACID* (*Aux/IAA*) genes during BnCN/PAN treatment (Fig. 8A, B; Additional file 11). A strong activation of central transcriptional regulators was observed, most notably *MONOPTEROS/ARF5* (*MP/ARF5*), which showed the highest expression induced by BnCN/PAN. This was accompanied by significant upregulation of multiple *ARFs* associated with developmental reprogramming and morphogenesis, including *ARF1*, *ARF3*, *ARF4*, *ARF6*, *ARF8*, *ARF10* and *ARF16*. In contrast, a pronounced repression of key negative regulators was detected, most prominently *ARF11* and *ARF20*. Among *Aux/IAA* genes 20/29 were differentially expressed, and most of them characterized with increased expression 13/20, especially *IAA1*, *IAA15*, *IAA20*, *IAA30,* and *IAA33*. To monitor auxin-responsive transcriptional activity during embryogenic induction, DR5::GUS derived signal was examined at days 1, 3, 5, 7, 10 and 15 of culture under BnCN/PAN treatment. Strong GUS activity was detected throughout the entire culture period and was broadly distributed across the explant (Fig. 8C). The signal was observed in all explant tissues as well as in developing somatic embryos, indicating sustained activation of auxin-responsive pathways during the acquisition and maintenance of embryogenic competence. Whereas, during pDR5::GUS seedling development in IZE explants not treated with BnCN/PAN, the GUS signal was restricted to the shoot apical meristem (SAM), the root apical meristem (RAM), and the cotyledon tips. Thus, we demonstrated that treatment of explants with BnCN/PAN activates auxin signaling.

**Figure 8.**
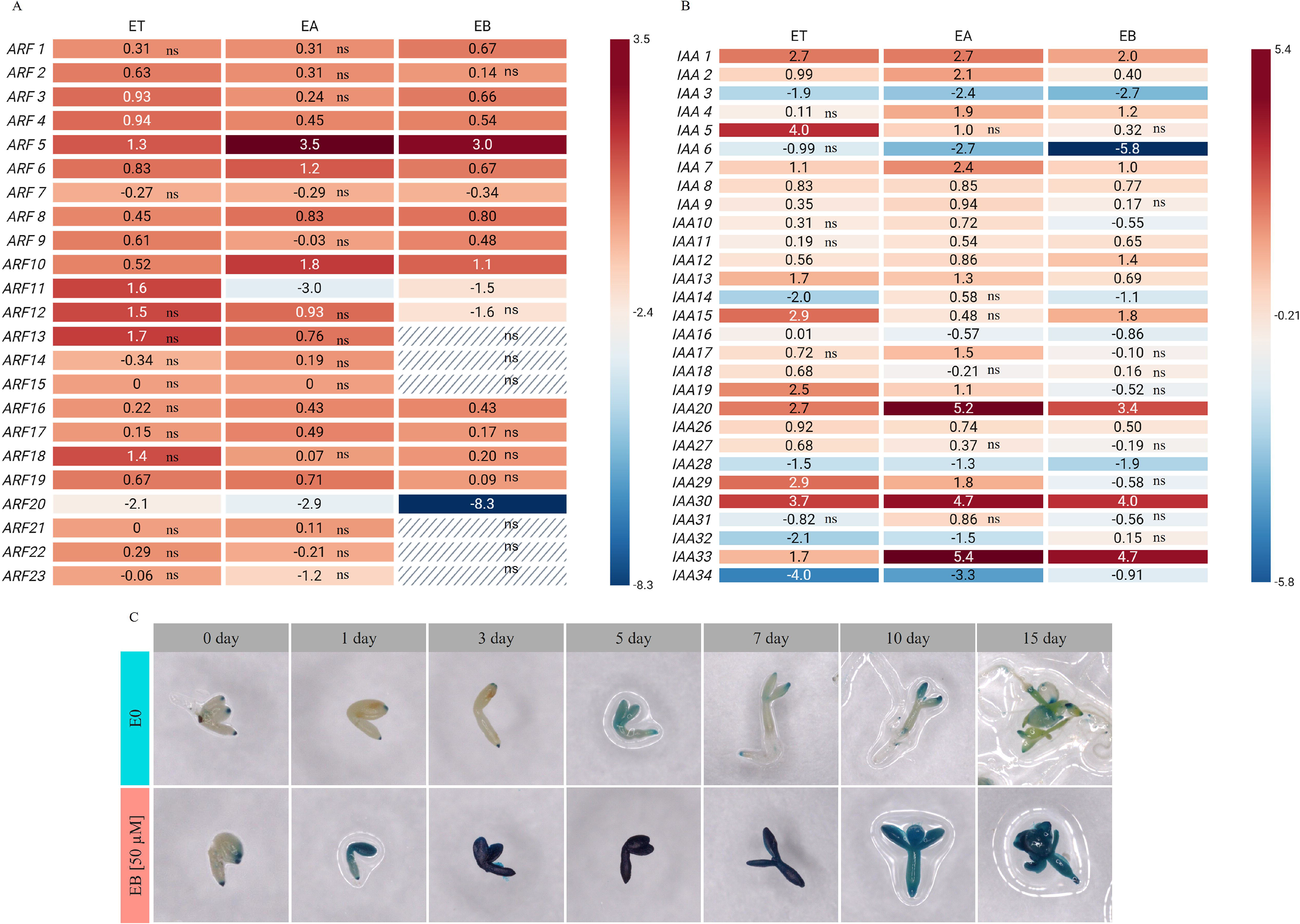
Induction of auxin signaling under BnCN/PAN treatment. Heatmaps showing differentially expressed genes under different SE-inducing treatments identified by RNA-seq, related to *ARF* (A) and *Aux/IAA* (B) genes family. Z-score scale bar from −5.8 (downregulation, in blue) to +5.4 (upregulation, in red). Spatiotemporal localization of the DR5::GUS signal, which was indicative of auxin accumulation and signalization in the explants cultured on the EB and E0 media (C).

## Discussion

Our results demonstrate that the BnCN/PAN treatment efficiently induces SE process in *Arabidopsis thaliana* immature zygotic embryos, with optimal concentrations (≈50 µM) yielding up to 40% SE efficiency and ∼1.5 embryos per explant, while higher doses are inhibitory or toxic. The potential NIT-derived product PAA also triggers SE, reaching nearly 100% efficiency at 40 µM, confirming that both BnCN/PAN and PAA activate the embryogenic program, although less efficiently than 2,4-D, most frequently used for SE induction (Wójcik et al., 2020). Genetic analysis shows partial redundancy of *NIT* genes, with only *nit1-3* affecting BnCN/PAN-induced SE process, while inhibition of NITs by heatin reduces embryogenic response under both BnCN and 2,4-D treatment. Transcriptomic analysis reveals strong reprogramming, including activation of cell cycle, hormone signaling, and auxin-related pathways, alongside repression of stress and secondary metabolism genes. A conserved core of 3 659 DEGs is shared across different SE inducers, indicating a common regulatory network. BnCN/PAN strongly activates auxin signaling and key SE regulators such as *LAFL*, *WOX*, and *PLT* genes.

### BnCN/PAN as a substrate for NIT-dependent PAA biosynthesis

We hypothesize that BnCN/PAN may itself act as a bioactive compound capable of inducing an embryogenic response, or, more plausibly, BnCN/PAN may serve as a substrate for the biosynthesis of the naturally occurring *Arabidopsis thaliana* auxin PAA (Urbancsok et al., 2018, Perez et al., 2023a, Günther et al., 2018, Zhao et al., 2026). The study demonstrates that BnCN/PAN functions as a key intermediate in the PAOx-derived PAA biosynthetic pathway. Isotopic labeling experiments revealed that phenylacetaldoxime (PAOx) is converted to BnCN/PAN *in vivo*. Furthermore, feeding experiments with ¹³C-labeled BnCN/PAN confirmed its subsequent conversion into PAA, as ¹³C-PAA was detected exclusively in treated seedlings, establishing BnCN/PAN as a direct precursor of PAA (Zhao et al., 2026). The conversion of BnCN/PAN into PAA is most likely mediated by NIT enzymes. Among seven putative nitrilase genes in the *Solanum lycopersicum* genome, *Solyc11g068730* (designated *SlNIT1*) clustered phylogenetically with functionally characterized NITRILASES and shared >60% amino acid identity with validated enzymes, whereas the remaining candidates showed substantially lower similarity. Recombinant *SlNIT1* expressed in *Escherichia coli* efficiently converted BnCN/PAN to PAA *in vitro*, while no product formation was observed in control reactions. Together, these findings demonstrate that SlNIT1 catalyzes the hydrolysis of BnCN/PAA to PAA and support a PAOx – BnCN/PAN -PAA biosynthetic route in tomato (Zhao et al., 2026), establishing NITRILASES as key enzymatic components of this auxin-related metabolic pathway.

### Use of PAA in plant *in vitro* cultures

The use of PAA for the induction of *in vitro* regeneration in plants is well documented, in contrast to BnCN/PAN. PAA is binding to TRANSPORT INHIBITOR RESPONSE 1 (TIR1) AUXIN SIGNALING F-BOX PROTEIN 5 (AFB5), however its affinity for nuclear auxin receptors is lower than that of IAA (Lee et al., 2014); PAA is generally considered less bioactive than IAA (Sugawara et al., 2015), and is not directionally transported (Ung et al., 2022); nevertheless, it displays a substantially higher, approximately 50% greater, dose-dependent capacity to induce somatic embryogenesis (this study) compared with IAA (Wójcikowska et al., 2025). It was reported previously that PAA enhances callus formation (Leuba and LeTourneau, 1990). Notably, there are scientific reports indicating that PAA can be applied to induce embryogenic processes i.e., somatic embryogenesis, androgenesis, gynogenesis, as well as shoot organogenesis (Table 1). The concentration of PAA used for the induction of somatic embryogenesis in *Pelargonium × hortorum* hypocotyl cultures, which resulted in efficient SE induction, ranged from 10–20 µM PAA. In our case, PAA concentrations in the range of 20-40 µM also resulted in the most efficient induction of SE process.

**Table 1.**
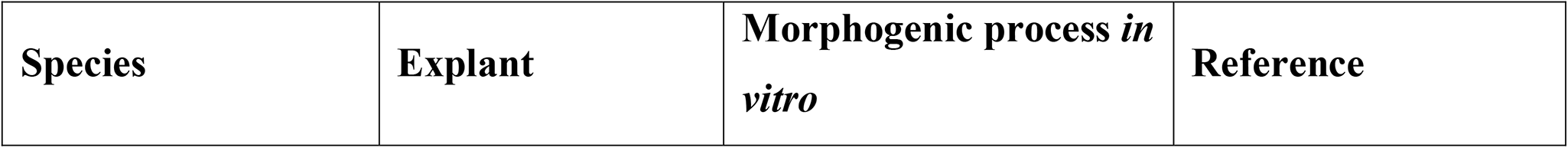
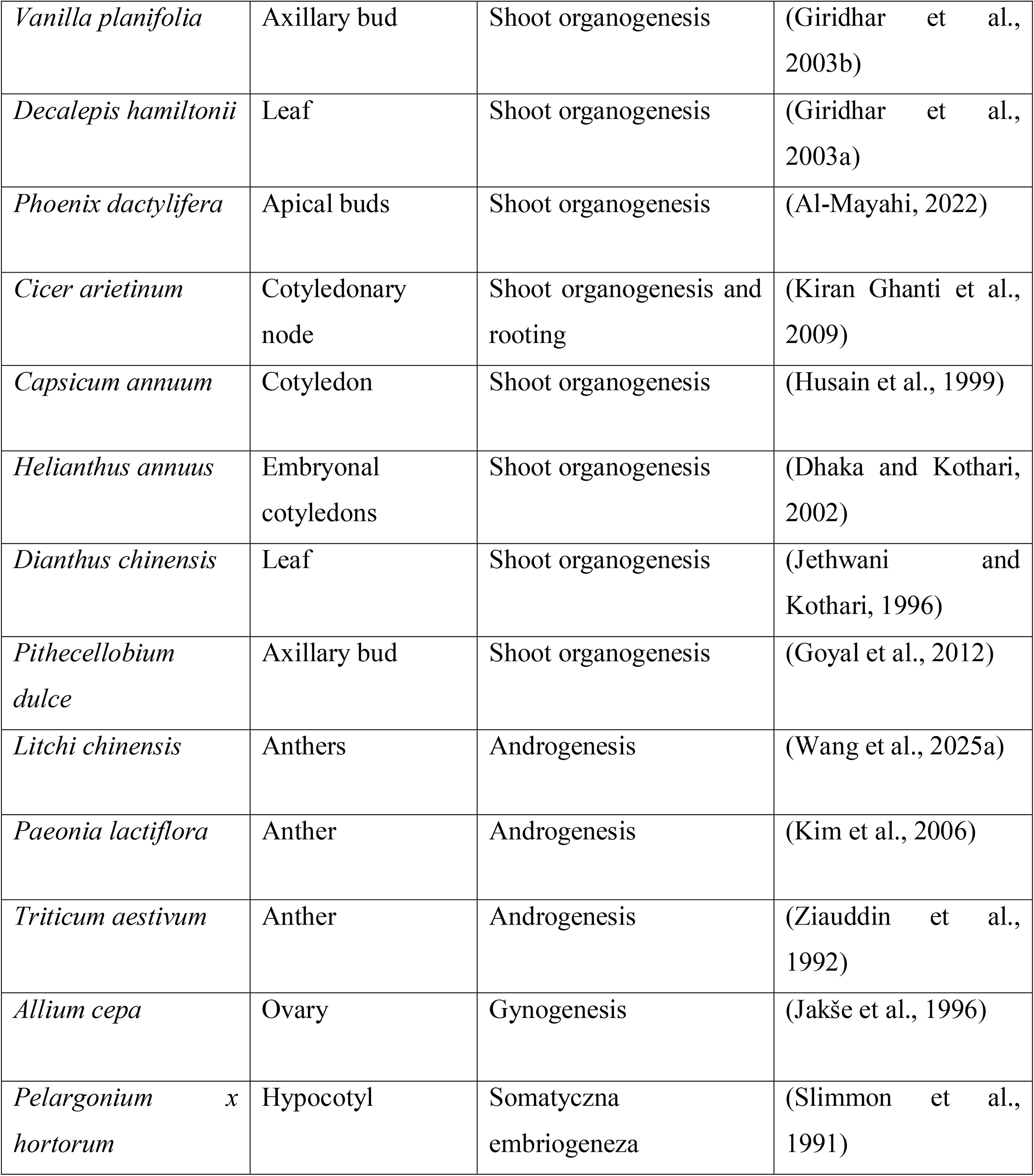

### The role of *NITs* in the process of somatic embryogenesis

Reports on the involvement of NITs in regenerative processes under *in vitro* culture conditions are scarce. It was noticed that, NITs are involved in process of embryogenic transition as inhibition of NIT activity using heatin (presented data) or mutations in the *NIT1*-*NIT4* genes results in a reduced embryogenic competence of the *Arabidopsis thaliana* explants (Wójcikowska et al., 2024). Furthermore, in *Arabidopsis thaliana lec1* mutant, unable to regenerate somatic embryo, the expression of *NIT1*, *NIT2*, and *NIT4* is highly decreased, which provides additional evidence supporting the involvement of *NITs* in the SE process (Wójcikowska et al., 2024). Additionally, the study demonstrated that LaMIR166a promotes somatic embryo maturation by enhancing *LaNIT* expression and endogenous auxin accumulation. Increased *LaNIT* transcript abundance was associated with elevated auxin signaling (*LaARF1* and *LaARF2*) and reduced embryo abnormalities, suggesting that nitrilase-mediated auxin biosynthesis is required for proper somatic embryo development in *Larix leptolepis*. However, among auxin only IAA level was analyzed, not PAA (Li et al., 2018).

### The effect of BnCN/PAN on auxin signaling

Why BnCN/PAN and PAA triggers the embryogenic programme remains largely unresolved. Using global transcriptome profiling, we demonstrated that BnCN/PAN activates the expression of genes associated with auxin signaling, thereby supporting earlier hypotheses and the limited evidence suggesting that BnCN/PAN-derived PAA is capable of activating auxin signaling pathways (Shimizu-Mitao and Kakimoto, 2014). We demonstrated that 30% of *ARF* genes were upregulated after BnCN/PAN treatment, including several *ARFs* previously reported to play key roles in regeneration, mostly SE process i.e., *ARF1*, *ARF3* (Wójcikowska and Gaj, 2017), *ARF4* (Zhang et al., 2021), *MP/ARF5* (Wójcikowska et al., 2025)*, ARF6*, *ARF8* (Su et al., 2016), *ARF10* and *ARF16* (Wójcik et al., 2017). Similarly, among *Aux/IAA* genes 45% of genes were characterized with increased expression under BnCN/PAN-dependent SE induction, especially *IAA1*, *IAA15*, *20*, *30,* and *33*. Aux/IAA proteins are key components of the nuclear auxin signaling pathway, where they act as transcriptional repressors under basal conditions and permit auxin-responsive gene activation following their rapid degradation. In recent years, extensive studies of Aux/IAA proteins have established their broad involvement in plant development, encompassing primary and lateral root formation, maintenance of apical dominance, gravitropic responses, the regulation of flower and fruit development, as well as stress response (Kumar et al., 2026). Moreover, *Aux/IAA* genes are involved in embryogenic transition in many species (Wójcik et al., 2020, Zhou et al., 2024, Wang et al., 2025b, Yang et al., 2025, Peng et al., 2022, Kang et al., 2021). Furthermore, it has been shown that several *Arabidopsis thaliana aux/iaa* mutants, including *iaa16, 29, 30,* and *31*, characterized with decreased capacity to SE process (Gliwicka et al., 2013). It was shown that seedlings exposure to 50 and 100 μM BnCN/PAN induced the expression of *IAA5, 12, 19, 29* (Urbancsok et al., 2018). This suggests that, depending on the tissue type, distinct subsets of *Aux/IAA* genes are differentially regulated by BnCN/PAN. In our transcriptomic data we identified also four *SAUR* (*SMALL AUXIN UP RNA*) genes (*SAUR8 50, 51, 54*) highly activated after BnCN/PAN treatment and embryogenic transition, similarly in other plant species *SAUR* were characterized with increased expression under SE induction (Zanin et al., 2022, Chen et al., 2023). *SAURs* belong to a large family of early auxin-responsive genes and are involved in regulating plant growth and many developmental processes (Chen et al., 2023). It was showed that *DgSAUR1* is a highly sensitive and specific transcriptional marker of PAA signaling. Exogenous PAA strongly induced *DgSAUR1* expression in *Datisca glomerata* roots, confirming the activation of auxin-responsive transcriptional programs (Salgado et al., 2025). A summarizing, this promoting effect of BnCN/PAN on SE-process could be attributed to BnCN/PAN-NITs-dependent PAA biosynthesis, which potentially regulates endogenous auxin signaling and enhances the morphogenesis of early somatic embryos.

### The effect of BnCN/PAN on expression of genes encoding embryogenic master regulators

Treatment of explants with BnCN/PAN leads to increased expression of genes encoding transcription factors (LAFL; PLT; WOXs) and EMB proteins that play key roles in somatic embryogenesis. The functions of these proteins (LAFL; PLT; WOXs, EMB) have been well documented and reviewed, demonstrating that loss-of-function mutations suppress, whereas overexpression frequently leads to activation of the embryogenic program in plants. Consequently, they are widely exploited in plant biotechnology to enhance regenerative capacity (Chen et al., 2024). Finally, we experimentally validated the hypothesis proposed by Wang et al. (2025a), demonstrating that the high frequency of shoot regeneration from PAA-treated somatic embryos suggests PAA direct involvement in somatic embryogenesis, potentially through modulation of key transcription factors such as *LEC*, *WUS*, and *PLT4*/*BABY BOOM* (*BBM*).

### The involvement of BnCN/PAN in plant *in vitro* regenerative processes

To the best of our knowledge, this study provides the first evidence that BnCN/PAN can induce embryogenic transition in *in vitro* cultures of immature zygotic embryos in plants, using *Arabidopsis thaliana*. To date, no studies have explored the use of BnCN/PAN in plant *in vitro* systems for the induction of other embryogenesis-related processes, including somatic embryogenesis, androgenesis, and gynogenesis. Similarly, its potential to induce organogenesis, such as shoot or root regeneration, remains unknown. Collectively, these findings position BnCN/PAN among the expanding group of organic growth-promoting compounds with the capacity to enhance plant regeneration *in vitro* (Hamdeni et al., 2022, Samiei et al., 2021). Obtained results open new research perspectives and prompt further questions regarding the mechanisms underlying BnCN/PAN-mediated regulation of regeneration. Can BnCN/PAN reduce genotype-dependent variation in the *in vitro* response of plant explants and enhance somatic regeneration in recalcitrant genotypes? What are the functional and metabolic relationships among BnCN/PAN, PAA, IAA, and other naturally occurring auxins during the induction of *in vitro* plant regeneration? Does BnCN/PAN act as a metabolic precursor of PAA in plant tissues, and is its effect on SE dependent only on NITRILASE activity?

## Materials and Methods

### Plant material

The Columbia (Col-0) genotype of *Arabidopsis thaliana* (L.) Heynh. Col-0, *pDR5::GUS* reporter line, insertional and chemical-induced *nit* mutant seeds were supplied by NASC (The Nottingham Arabidopsis Stock Centre) (Additional file 12).

### Plant growth and in vitro culture conditions

Seed-derived plants were grown in Jiffy*-*7 (Jiffy, Norway) peat pots in a “walk-in” type phytotron under controlled conditions at 22°C under a 16 h photoperiod of 100 µM m^-2^ s^-1^ white, fluorescent light. The *in vitro*-grown plant material, grown in sterile conditions, was kept at 23°C under a 16/8 h (light/dark) photoperiod and 40 µM m^-2^ s^-1^ white fluorescent light.

### In vitro culture of explants and SE induction

Immature zygotic embryos (IZEs) of *Arabidopsis thaliana* at the cotyledonary stage were used as explants for *in vitro* culture. The explants were isolated from siliques and cultured following the standard protocol for SE induction (Gaj, 2001). A basal E0 medium contained: 3.2 g L^-1^ of B5 micro and macro-elements (Duchefa Biochemie; #G0210)(Gamborg et al., 1968), 20 g L^-1^ sucrose and 8 g L^-l^ agar, pH 5.8. The SE induction media contained: 5 μM 2,4-D () used as a control or 25-600 μM (25, 50, 75, 100, 200, 300, 400, 500, 600 μM) BnCN/PAN (Sigma Aldrich; #140-29-4). In addition, E0 medium supplemented with PAA (Serva; #32106) at a concentration of 5, 10, 20, 30, 40, 50, 75, 100, 125, 150 μM was applied for SE induction. Heatin (Chembridge **#**5713980) dissolved in DMSO was used to inhibit NITs in EB, EA media at concentration of 15 μM. The SE capacity was evaluated in three-week-old cultures. Two parameters, i.e., SE efficiency, calculated as the frequency of the explants that produced somatic embryos, and SE productivity, calculated as the average number of somatic embryos per explant, were used to analyze the SE capacity of the studied lines. Thirty explants, with at least three replicates, were evaluated for each culture combination.

### GUS detection

Explants of the *pDR5::GUS* line cultured on the medium supplemented with 50.0 μM BnCN for 0, 1, 3, 5, 7, 10, 15 days were sampled and incubated in a GUS assay solution at 37°C for 12 hours (Jefferson et al., 1987). Pigments from tissue were removed with 95% ethanol. The experiments were repeated twice, and at least 30 somatic embryos were subjected to GUS signal detection in each repetition. The GUS staining pattern was analyzed using a stereomicroscope KEYENCE VHX-97OF.

### RNA isolation, library preparation, and sequencing

A NucleoSpin RNA PlantKit (Macherey-Nagel) was used to isolate total RNA from the IZE explants induced on medium containing 50 μM of BnCN/PAN (EB) for 10 days and control medium without SE-inducing agent (E0). The freshly isolated tissues were wiped in frozen mortars. Around 100 (10^th^ day) explants were used for RNA isolation per repetition. RNA isolation, library preparation, and sequencing were produced in four biological replicates. The concentration and purity of RNA samples were evaluated with an ND-1000 spectrophotometer (NanoDrop Technologies).

### mRNA library construction

Messenger RNA was purified from total RNA using poly-T oligo-attached magnetic beads. After fragmentation, the first strand cDNA was synthesized using random hexamer primers. Then the second strand cDNA was synthesized using dUTP, instead of dTTP. The directional library was ready after end repair, A-tailing, adapter ligation, size selection, USER enzyme digestion, amplification, and purification. The library was checked with Qubit and real-time PCR for quantification and bioanalyzer for size distribution detection. After library quality control, different libraries were pooled based on the effective concentration and targeted data amount, then subjected to Illumina sequencing. The basic principle of sequencing is sequencing by Synthesis, where fluorescently labeled dNTPs, DNA polymerase, and adapter primers are added to the sequencing flow cell for amplification. As each sequencing cluster extends its complementary strand, the addition of each fluorescently labeled dNTP releases a corresponding fluorescence signal. The sequencer captures these fluorescence signals and converts them into sequencing peaks through computer software, thereby obtaining the sequence information of the target fragment. Only libraries meeting the quality criteria were pooled based on concentration and sequencing requirements and subjected to high-throughput sequencing on an Illumina platform, following Novogene’s standard workflow.

### Sequencing quality control

Raw data (raw reads) of fastq format were firstly processed through fastp software. In this step, clean data (clean reads) were obtained by removing reads containing adapter, reads containing ploy-N and low quality reads from raw data. At the same time, Q20, Q30 and GC content the clean data were calculated. All the downstream analyses were based on the clean data with high quality.

### Read alignment to the reference genome

The *Arabidopsis thaliana* reference genome and annotation files (TAIR10 version) were downloaded from Ensembl Plants (https://plants.ensembl.org). Reference genome and gene model annotation files were downloaded from genome website. Use HISAT2 (2.2.1) to build the index of the reference genome, and use HISAT2 to align paired-end clean reads to the reference genome. HISAT2 can use the gene model annotation file to create splice-aware alignments, providing better alignment accuracy compared to other non-splice alignment tools.

### Gene expression quantification

The featureCounts (2.0.6) was used to count the reads numbers mapped to each gene. And then FPKM of each gene was calculated based on the length of the gene and reads count mapped to this gene. FPKM, expected number of Fragments Per Kilobase of transcript sequence per Millions base pairs sequenced, considers the effect of sequencing depth and gene length for the reads count at the same time, and is currently the most commonly used method for estimating gene expression levels..

### Differential expression analysis

For DESeq2 with biological replicates: Differential expression analysis for two conditions/groups was performed using the DESeq2 R package (1.42.0). DESeq2 provides statistical programs for determining differential expression in digital gene expression data using models based on negative binomial distribution. The resulting P-value is adjusted using the Benjamini and Hochberg’s methods to control the error discovery rate. The threshold of significant differential expression: padj <= 0.05, |log2(foldchange)| >= 1. For edgeR without biological replicates: Prior to differential gene expression analysis, for each sequencing library, read counts were adjusted using the edgeR R package (4.0.16) by scaling normalization factors to eliminate differences in sequencing depth between samples, followed by differential expression analysis. The resulting P value is adjusted using the Benjamini and Hochberg’s methods to control the error discovery rate. The threshold of significant differential expression: padj <= 0.05, |log2(foldchange)| >= 1.

### GO and KEGG enrichment analysis of differentially expressed genes

Gene Ontology (GO) enrichment analysis of differentially expressed genes was implemented by the clusterProfiler (4.8.1), in which gene length bias was corrected. GO terms with corrected P-value less than 0.05 were considered significantly enriched by differential expressed genes. KEGG is a database resource for understanding high-level functions and utilities of the biological system, such as the cell, the organism and the ecosystem, from molecular-level information, especially large-scale molecular datasets generated by genome sequencing and other high-through put experimental technologies (http://www.genome.jp/kegg/). We used clusterProfiler R package to test the statistical enrichment of differential expression genes in KEGG pathways. Gene Set Enrichment Analysis (GSEA) is a computational approach to determine if a predefined Gene Set can show a significant consistent difference between two biological states. The genes were ranked according to the degree of differential expression in the two samples, and then the predefined Gene Set were tested to see if they were enriched at the top or bottom of the list. Gene set enrichment analysis can include subtle expression changes. We use the local version of the GSEA analysis tool http://www.broadinstitute.org/gsea/index.jsp, GO, KEGG data set were used for GSEA independently. PPI analysis of differentially expressed genes was based on the STRING database, which contains known and predicted protein-protein interactions. For species present in the database, we construct the network by extracting the target gene list from the database. Otherwise, we use diamond (version 0.9.13) to align the target gene sequences with selected reference protein sequences, and then establish the network based on the known interactions of the selected reference species.

### Statistical analysis and data visualization

The Student’s t-test (p < 0.05) or two-way ANOVA analysis (p < 0.05) followed by Tukey’s HSD test (p < 0.05) were used to determine any significant differences between the compared combinations. The graphs show the means with the standard deviations (SD). BioRender was used to create the graphical abstract and figures.

## The following materials are available in the online version of this article. Supplementary Data

**Additional file S1.** List of differentially expressed genes under BnCN/PAN treatment during embryogenic transition.

**Additional file S2.** Expression profile of all *Arabidopsis thaliana* genes under BnCN/PAN treatment during embryogenic transition.

**Additional file S3.** List of downregulated DEGs under BnCN/PAN treatment during embryogenic transition.

**Additional file S4.** List of upregulated DEGs under BnCN/PAN treatment during embryogenic transition.

**Additional file S5.** List of DEGs under BnCN/PAN treatment during embryogenic transition in sub cluster 1.

**Additional file S6.** List of DEGs under BnCN/PAN treatment during embryogenic transition in sub cluster 2.

**Additional file S7.** List of DEGs under BnCN/PAN treatment during embryogenic transition in sub cluster 3.

**Additional file S8.** List of DEGs under BnCN/PAN treatment during embryogenic transition in sub cluster 4.

**Additional file S9.** List of shared and treatment-specific DEGs after BnCN/PAN (EB medium), 2,4-D (EA medium) and TSA (ET medium) IZE treatment causing embryogenic transition.

**Additional file S10.** Expression profiles of genes encoding ARF and Aux/IAA transcription factors acting as under BnCN/PAN, 2,4-D, and TSA treatments leading to somatic embryogenesis induction.

**Additional file S11.** Expression profiles of genes encoding NITs, transcription factors acting as master regulators of somatic embryogenesis (SE-MR), and EMB proteins under BnCN/PAN, 2,4-D, and TSA treatments leading to somatic embryogenesis induction.

**Additional file S12.** Insertional and chemical-induced *nit* mutant seeds with NASC ID.

## Acknowledgments

We acknowledge Jacek Pietrakowski and Katarzyna Konopka for technical assistance.

## Data and resource availability

Raw images and source data are provided in the Zenodo repository (10.5281/zenodo.21352216). The RNA-seq raw data presented in this publication have been deposited in NCBI’s Gene Expression Omnibus (GEO) and are accessible through GEO Series accession number GSE339067. Any additional information required to re-analyze the data reported in this paper is available from the lead contact upon request.

## Author contributions

B.W. designed and supervised the project. B.W., J.F., conducted the experiments and analyzed the data. B.W. wrote the manuscript. B.W., M.M. provided funding. All authors discussed the results and approved the manuscript.

